# Tuning TPO-R signaling to influence hematopoietic stem cell differentiation and inhibit essential thrombocythemia

**DOI:** 10.1101/2020.09.23.290593

**Authors:** Lu Cui, Ignacio Moraga, Tristan Lerbs, Camille Van Neste, Stephan Wilmes, Naotaka Tsutsumi, Aaron Claudius Trotman-Grant, Milica Gakovic, Sarah Andrews, Jason Gotlib, Spyridon Darmanis, Martin Enge, Stephen Quake, Ian S. Hitchcock, Jacob Piehler, K. Christopher Garcia, Gerlinde Wernig

## Abstract

Thrombopoietin (TPO) and the TPO-receptor (TPO-R, or c-MPL)) are essential for hematopoietic stem cell (HSC) maintenance and megakaryocyte differentiation. Agents that can modulate TPO-R signaling are highly desirable, both experimentally and clinically. We have developed a series of surrogate protein-ligands for TPO-R, in the form of diabodies, that homodimerize the TPO-R on the cell surface in different geometries, in effect ‘tuning’ downstream signaling responses. These surrogate ligands exhibit diverse pharmacological properties, inducing graded signaling outputs, from full to partial TPO agonism and antagonism, thus decoupling the dual functions of TPO/TPO-R. Using scRNA sequencing and HSC self-renewal assays we find that partial agonistic diabodies preserved the stem-like properties of cultured HSCs, but also blocked oncogenic colony formation in Essential Thrombocythemia (ET) through inverse agonism. Our data suggest that dampening downstream TPO signaling is a powerful approach not only for HSC preservation in culture, but also for inhibiting oncogenic signaling through the TPO-R.

**Significance Statement:** The TPO cytokine, which signals through its receptor c-MPL (or TPO-R), is essential for megakaryocyte differentiation and maintenance of hematopoietic stem cells (HSCs). Its signaling is deregulated in Essential Thrombocythemia (ET). Here, we engineered diabodies (DBs) against the TPO-R as surrogate TPO ligands to manipulate TPO-R signaling, from full to partial to antagonism, thus decoupling the dual functions of TPO/TPO-R (i.e, HSC maintenance versus megakaryopoiesis). We subsequently discovered that partial agonistic DBs, by reducing the strength of the TPO-R signal, not only preserved HSCs in culture, but also blocked oncogenic signaling in ET. This finding has the potential to improve HSC cultures for transplants, as well as serve as a unique therapeutic approach for ET.

## Introduction

Since the landmark cloning of thrombopoietin (TPO) (1–4) and its receptor MPL (TPO-R) (5–7), there has been interest in characterizing the role of the TPO/TPO-R interaction in normal and pathological hematopoiesis. TPO signaling plays key roles in regulating megakaryopoiesis and platelet production (8, 9) and in the support of hematopoietic stem cell (HSC) maintenance and self-renewal (10–15). Consequently, aberrant TPO/TPO-R signaling can lead to severe hematological disorders including myeloproliferative neoplasms (MPNs) (16, 17) and bone marrow failure syndromes (18, 19).

Despite initial hurdles with first generation recombinant thrombopoietin, which stimulated autoantibody formation resulting in cytopenia, several second-generation peptide/small-molecule TPO mimetics are now approved for immune thrombocytopenia and hepatitis C-associated thrombocytopenia and under evaluation for a wide range of other thrombocytopenic disorders, including those associated with chemotherapy and bone-marrow failure syndromes. Side effects of TPO mimetics include thrombocytosis, possibly thrombosis, bone marrow reticulin fibrosis, and rebound thrombocytopenia (20). Thus, fine-tuning this therapeutically important signaling axis remains an important goal.

TPO is a prototypical four-helix bundle cytokine(21), and engages TPO-R using the canonical “growth hormone” mechanistic and structural paradigm (22, 23). TPO dimerizes TPO-R and induces downstream signaling through the JAK/STAT pathway, principally activating JAK2 and STAT5 (24). However, few studies have addressed how TPO-R dimerization dynamics and geometry, in response to TPO, relates to signaling output, and whether TPO-R signaling could be manipulated in therapeutically useful ways that are not exhibited by the natural TPO ligand. This is important given that the activities of the natural TPO molecule are clinically limited (25). Although synthetic small molecule TPO-R agonists have been developed (21), as well as agonist antibodies (26), they appear to phenocopy TPO-induced signaling, and lack the element of fine-control of agonist output.

Two particularly relevant studies have hinted at the potential promise of diversifying TPO pharmacology. In one study, the authors showed that artificial chimeric TPO-R fusion proteins, when forced into different symmetrical dimeric receptor orientations, differentially promoted megakaryocyte differentiation, cell adhesion, and myeloproliferative and myelodysplasia phenotypes in vivo (27). This intriguing result raised the possibility that engineered ligands that could bind to unmodified TPO-R on natural cells and alter dimer geometry, could potentially induce different functional outcomes. Following this observation, we demonstrated that a dimeric antibody format known as diabodies (DBs), had the capacity to bind to different epitopes on the EPO-R, a highly-related class I cytokine receptor, and induce different dimerization geometries, modulating signal strength (28). These DBs elicited a range of signaling pharmacology, from full to partial agonism. Given that TPO-R and EPO-R both form JAK2-associated homodimeric signaling complexes on the cell surface, and exhibit highly similar signaling signatures, these studies motivated us to ask if we could manipulate TPO-R signaling with engineered surrogate ligands.

In the present study, we manipulate TPO-R signal strength, borrowing a strategy commonly used for GPCRs that has been rarely applied to cytokine receptors. We used DBs as ‘surrogate’ TPO-R ligands that bind to different epitopes on the TPO-R extracellular domain (ECD), and as a result, alter the TPO-R inter-dimer distance and geometry which results in a perturbed, and fine-tuned, TPO-R downstream signal. We evaluated the activities of these TPO-R DBs by measuring the level of TPO-R dimerization and signaling downstream and exploring their effects on HSC gene expression and differentiation. We also transcriptionally single-cell profiled the effects the DBs and TPO had on proliferation and differentiation of FACS-purified human HSC after ex vivo expansion (29, 30). In addition, the cytokine TPO is required in all cell culture media to expand HSC ex vivo for cell therapy (31). Therefore, we evaluated whether substituting TPO with our surrogate agonist TPO-R DBs could influence cell-fate decisions and increase the HSC number for allogenic hematopoietic stem-cell transplantation. Several of the partial agonist TPO-R DB molecules control HSC maintenance versus differentiation under ex vivo expansion and differentiation conditions, thus pointing to tuning of TPO-R signal strength as a major determinant of HSC homeostasis. Collectively these studies show that DBs can split the dual function of TPO/TPO-R (megakaryopoiesis and HSC maintenance) and that the calibration and tuning of TPO-R can exert differential effects on HSC fate decisions.

## Results

### Diabodies Induce Different Degrees of Agonistic Activity

As a proxy for the natural TPO ligand, we used diabodies (DBs), which have highly engineerable scaffolds and the capacity to dimerize their targets because they contain two binding sites. DBs are a dimeric, bivalent antibody format generated through a tandem arrangement of two single-chain scFv’s connected by a short linker. This format results in a dimeric molecule in which the binding sites are closer together and more rigidly connected than a typical IgG-like antibody where two Fabs are linked by an Fc. The closer proximity of the two binding sites in the DBs increases chances for productive TPO-R dimerization and signaling. Previously, a full agonist DB against TPO-R was reported, showing that a DB can in principle dimerize TPO-R in a productive signaling geometry (26). We generated three versions of DBs using sequences of anti-TPO-R DBs, as described in methods (26, 28, 30). Each of these DBs has different sequences of CDR loops, and therefore likely bind to different epitopes on TPO-R, thus dimerizing TPO-R in different geometries. We evaluated binding of the three DBs AK111, 113, 119 to the TPO-R receptor ectodomain (SD1-2) displayed on the surface of yeast, and found that all DBs bound with comparable efficiencies to TPO-R (**Fig. 1*A***), with DBs AK113 and AK111 binding having higher affinity. We next explored whether these DBs were functional in inducing TPO-R signaling. Stimulation of Ba/F3-MPL cells with TPO or the respective DBs induced TPO-R receptor phosphorylation across conditions, with levels significantly higher with TPO or AK119 than AK113 and AK111 treatment (**Fig. 1*B***). Upon TPO stimulation, TPO-R is phosphorylated at various tyrosine residues on its intracellular domain. To better understand the signaling properties of the different DBs and distinguish between complete and partial agonistic activity, we investigated the levels of receptor phosphorylation in three stable cell lines expressing TPO-R mutants where Tyr residues (all of which are phosphorylated upon TPO binding) were replaced with Phe (Y591F, Y625F and Y630F) (**Fig. 1*C***). While Y591F has been shown to be a negative regulator of TPO signaling and dispensable for proliferation (32), Y625F is essential for it. All three mutant cell lines showed high levels of phosphorylation with AK119 treatment (equal or higher than TPO). In contrast, phosphorylation was disrupted after AK111 treatment in all mutant cells suggesting partial agonism. Similarly, AK113 exhibited partial agonistic activities with a lesser phospho-TPO-R:TPO-R ratio than TPO or AK119 in wildtype TPO-R, but less in mutated TPO-R (**Fig. 1*D***). DBs AK119 and 113 show very similar responses to TPO. Indeed, a positive response is shown using doses in the nano-micromolar concentration range, whereas DB AK119 presents a very strong response within picomolar doses. Response saturation is very similar between DB AK 119 and TPO while saturation is observed with lower doses for DB AK111 and 113 (**Fig. 1*E***). This trend was further confirmed when we studied STAT5 phosphorylation (**Fig. 1*F***) and cell proliferation (**Fig. 1*G***) induced by the three DBs in UT-7 TPO-R cells.

**Fig. 1.**
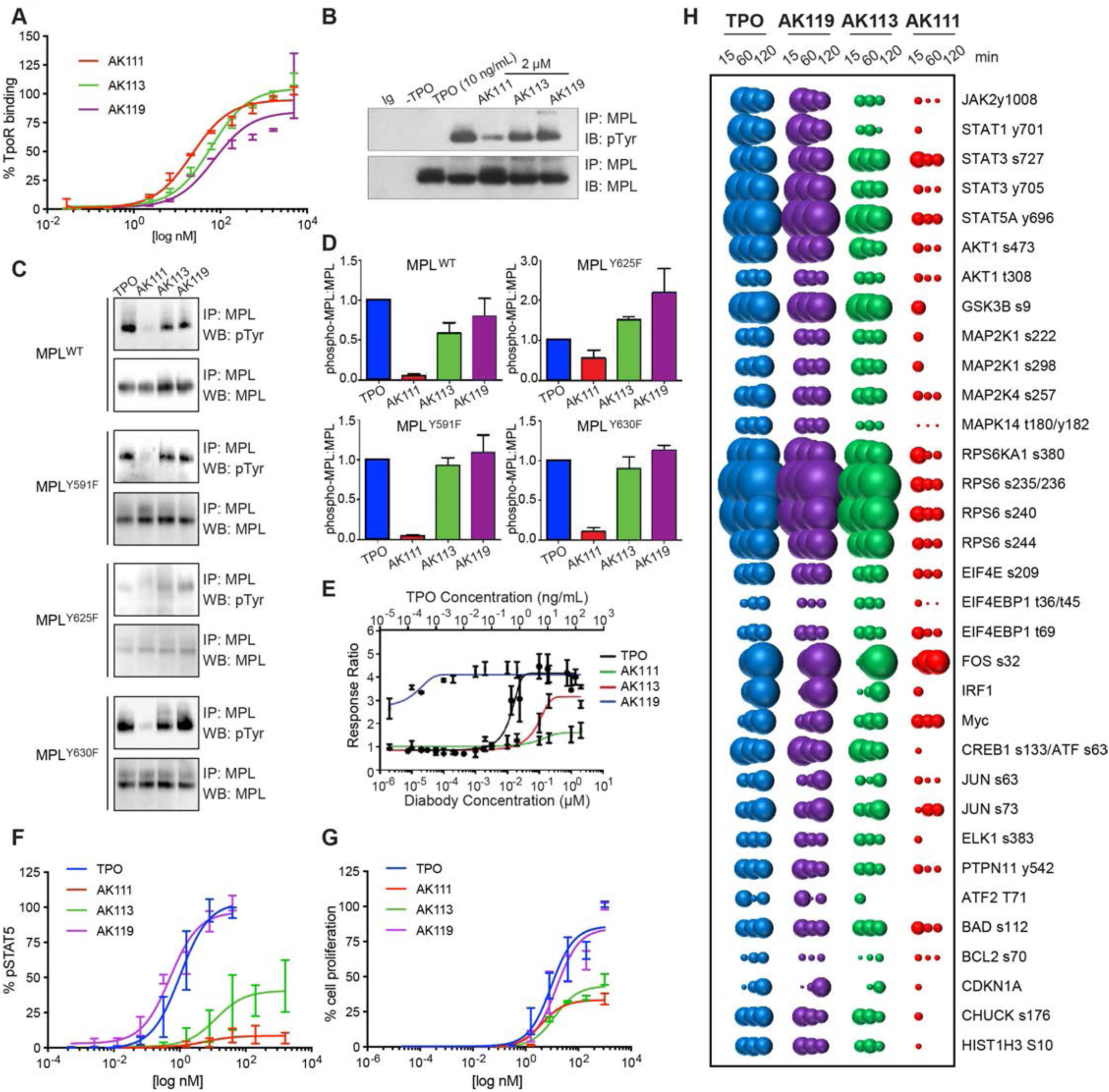
Diabodies induce different degrees of agonism activity. (*A*) Levels of TPO-R (MPL) binding (data are from independent replicates mean ± SD) and (*B*) levels of MPL phosphorylation promoted by DBs at the indicated dosages. (*C*) Study of the level of phosphorylation promoted by different DBs in different phospho mutants of MPL. (*D*) Ratio between the non-phosphorylated and phosphorylated MPL forms induced by different DBs. Data are from independent replicates mean ± SD. (*E*) Response ratio between low dose of TPO with high dose of TPO and DBs. Data are from independent replicates mean ± SD. (*F*) PhosphoSTAT5 dose-response experiments performed in UT-7 TPO-R cells stimulated with TPO or AK111-113-119 for 15 minutes. Data mean ± SD are from independent experiments. (*G*) UT-7 TPO-R proliferation in response to TPO or the three DBs for 5 days. (Data mean ± SD are from independent replicates). (*H*) Diabodies induce graded signaling strengths. Bubble plot representation of the signaling pathways activated by TPO and the DBs at the indicated times in UT-7-TPO-R cells. The size of the bubbles represents the intensity of the signal activated.

### Diabodies Induce Graded Signaling Strengths

TPO not only induces the activation of STAT5/STAT3/STAT1 but, in addition, it activates non-STAT pathways such as the MAPK and PI3K that further modify TPO-mediated responses (**Fig. 1*H* and S1**). Previously we showed for EPO-R that alterations on the ligand-receptor binding topology could lead to biased signaling (28), thus we asked whether the same principle would apply to the TPO-R system, and whether TPO-R DB were engaging these pathways as well. To test this, UT-7 TPO-R cells were stimulated with saturating concentrations of TPO or the three TPO-R DBs for the indicated times and the levels of over 133 molecules were analyzed via phospho-specific antibodies and flow cytometry. We detected activation of 33 different signaling molecules by TPO and the DBs (**Fig. 1*H***). These molecules included members of the STAT family (STAT1, STAT3 and STAT5), MAP kinase family (MEK, p38) and PI3K family (AKT, RSK1 and RPS6). We also found upregulation of known TPO-induced transcription factors such as IRF1 and c-MYC. In agreement with our previous results, the activities of the three DBs ranged from full agonism for AK119 to partial agonism for AK113 and weak agonism for AK111. Interestingly, not all 33 molecules were activated to the same extent by the DBs (**Fig. 1*H***). When the activation levels induced by the three DBs after 15 minutes of stimulation were normalized to those induced by TPO, we observed that while TPO and AK119 produced very similar signaling signatures, AK113 elicited a graded response, with some of the pathways more affected than others. For example, AK113 elicited almost full agonism for GSK3B and RPS6 activation but very weak agonism for STAT1 and STAT3 (**Fig. S1*A*)**. When the signaling output elicited by the EPO-R DB, the TPO-R DB, EPO and TPO were analyzed via principal component analysis, we noticed that AK113 elicited a signaling output in between full and minimal agonism and was closer to EPO and DA5 (**Fig. S1*B***). Overall our combined signaling data further confirm our previous observations that alterations on the ligand-receptor binding architecture results in fine-tuning of the signaling outputs delivered by cytokine receptors.

### Diabodies Dimerize TPO-R at the Surface of Living Cells

An important mechanistic question is whether the DB/TPO-R complexes on the cell surface are indeed homodimers or higher-order species resulting from clustering of TPO-R dimers. To test this, we employed dual-color single molecule total internal reflection fluorescence (TIRF) microscopy to quantify dimerization of TPO-R by different DBs in the plasma membrane of living cells. Selective and efficient labeling of TPO-R extracellularly fused to a non-fluorescent, monomeric EGFP (mXFP) was achieved via anti-GFP nanobodies conjugated with Rho11 and Dy647, respectively (**Fig. 2*A***). TPO-R dimerization was quantified by co-localization and co-tracking analysis of individual molecules with subdiffractional resolution as schematically depicted in **Fig. 2*A*** (Movie supplement 1 and **Fig. 2*B***). In the absence of stimulation, no co-trajectories were observed. Upon addition of TPO or DBs, however, co-locomoting TPO-R were clearly discerned, though with different efficiencies (Movie supplement 1-3 and **Fig. 2*B-C***). Relative dimerization levels observed for different DBs are summarized in **Fig. 2*D***. For DB AK119, similar, yet slightly increased dimerization as compared to TPO was observed. By contrast, dimerization by DB AK111 and 113 were significantly lower, though above the untreated condition. Since HeLa cell express low levels of JAK2, we co-expressed TPO-R with JAK2 fused to mEGFP (JAK2-mEGFP). In JAK2-mEGFP expressing cells, significantly increased TPO-R dimerization by DB AK119 was obtained, similarly to TPO. These higher dimerization levels may be ascribed to productive interactions of JAK2 homomers within the receptor complex. Interestingly, the addition of JAK2-mEGFP did not significantly increase TPO-R dimerization by DB AK111 and 113. This observation suggests that these DBs may form TPO-R dimers with a strongly altered geometry that do not allow such productive interactions between TPO-R-bound JAK2 molecules, which may explain their altered activity profiles.

**Fig. 2.**
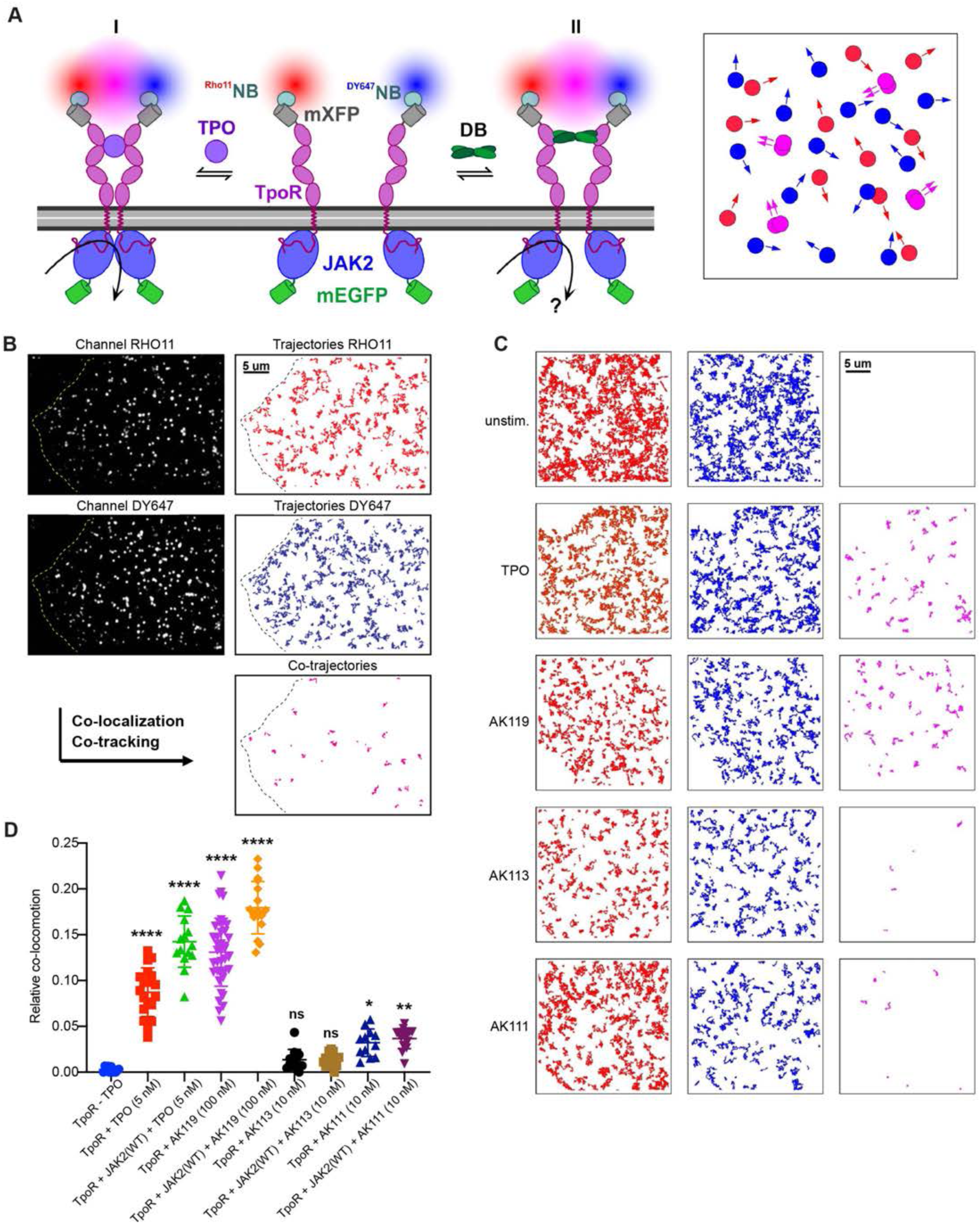
Diabodies dimerize TPO receptor at the surface of living cells. (*A*) Cell-surface labeling of TPO-R fused to a non-fluorescent mEGFP (mXFP) using dye-labeled (Rho11 and Dy647) anti-GFP nanobodies (NB). Co-expression of JAK2-mEGFP can be identified at single-cell level. Dimerization by TPO (I) and by DBs (II) is quantified by dual-color single-molecule co-tracking as schematically depicted on the right. (*B*) Representative trajectories (from 150 frames acquired in 4.8 s) of individual RHO11-labeled (red) and DY647-labeled (blue) TPO-R and co-trajectories (magenta) observed in a TPO-stimulated cell (cell boundaries are outlined by a dashed line). Scale bar: 5 µm. (*C*) Trajectories and co-trajectories observed in unstimulated cells and TPO-, AK119-, AK113- and AK111-treated conditions. Scale bars: 5 µm. (*D*) Changes in TPO- and DB-induced TPO-R dimerization upon co-expression of JAK2-mEGFP. Each data point represents the relative number of co-trajectories observed in individual cells. Data are expressed as mean ± SD and analyzed by Ordinary one-way ANOVA, * *P* < 0.05; ** *P* < 0.01; **** *P* < 0.0001, ns: non-significant.

### Diabodies Decouple the Dual Function of TPO/TPO-R during Megakaryocyte Differentiation and Block Oncogenic Signaling through the TPO-R

TPO plays a central role in the differentiation and maturation of megakaryocytes. Thus, we investigated the effects of TPO-R DBs in megakaryocyte differentiation. UT-7 cells have the potential to differentiate into megakaryocytes with TPO and to erythrocytes with EPO. To test the effects of DBs in this system, we incubated UT-7 cells with EPO, TPO or the partial or full agonistic TPO-R DBs for 24 days and then quantified megakaryocytic or erythrocytic differentiation in the cultures. AK113 stimulation resulted in weak upregulation of megakaryocytic genes and downregulation of erythroid genes, a physiologic process required during megakaryocytic differentiation (**Fig. S3**). Next, we analyzed the ability of the three TPO-R DBs to elicit megakaryocyte differentiation in liquid culture of freshly isolated human CD34+ HSCs from bone marrow cells in comparison to TPO. We also isolated megakaryocyte-erythroid progenitors (MEPs) by flow cytometry using the indicated markers (LIN-CD34+CD38+CD135-CD45RA-) and assessed colony-forming potential of the three DBs in direct comparison to TPO. We noticed that partial agonists AK111 and AK113 did not support effective megakaryocytic differentiation (**Fig. 3*A-C***) nor colony formation. Despite DBs AK111 and AK113 producing cells expressing megakaryocytic markers CD41 and CD42b, both resulted in significantly decreased numbers of polyploid cells in liquid and semisolid culture conditions, and were comparable in their low megakaryocyte differentiation potential to the low SCF (20 ng/ml) condition. In contrast, the full agonistic DB AK119 effectively promoted megakaryocytic differentiation and megakaryocytic colony formation to an even greater extent than TPO (**Fig. 3*A-C***). We also noted no significant contribution of TPO DBs to multipotent progenitors (MPP: lineage-CD34+CD38-CD90-CD45RA-), megakaryocyte-erythroid progenitors (MEP:lineage-CD34+CD38+CD135-CD45RA-), and granulocyte-macrophage progenitors (GMP: lineage-CD34+CD38+CD135+CD45RA+), and erythroid precursors, each accounting for <10% of cells both in liquid and semisolid culture conditions (**Fig. S4-6**).

**Fig. 3.**
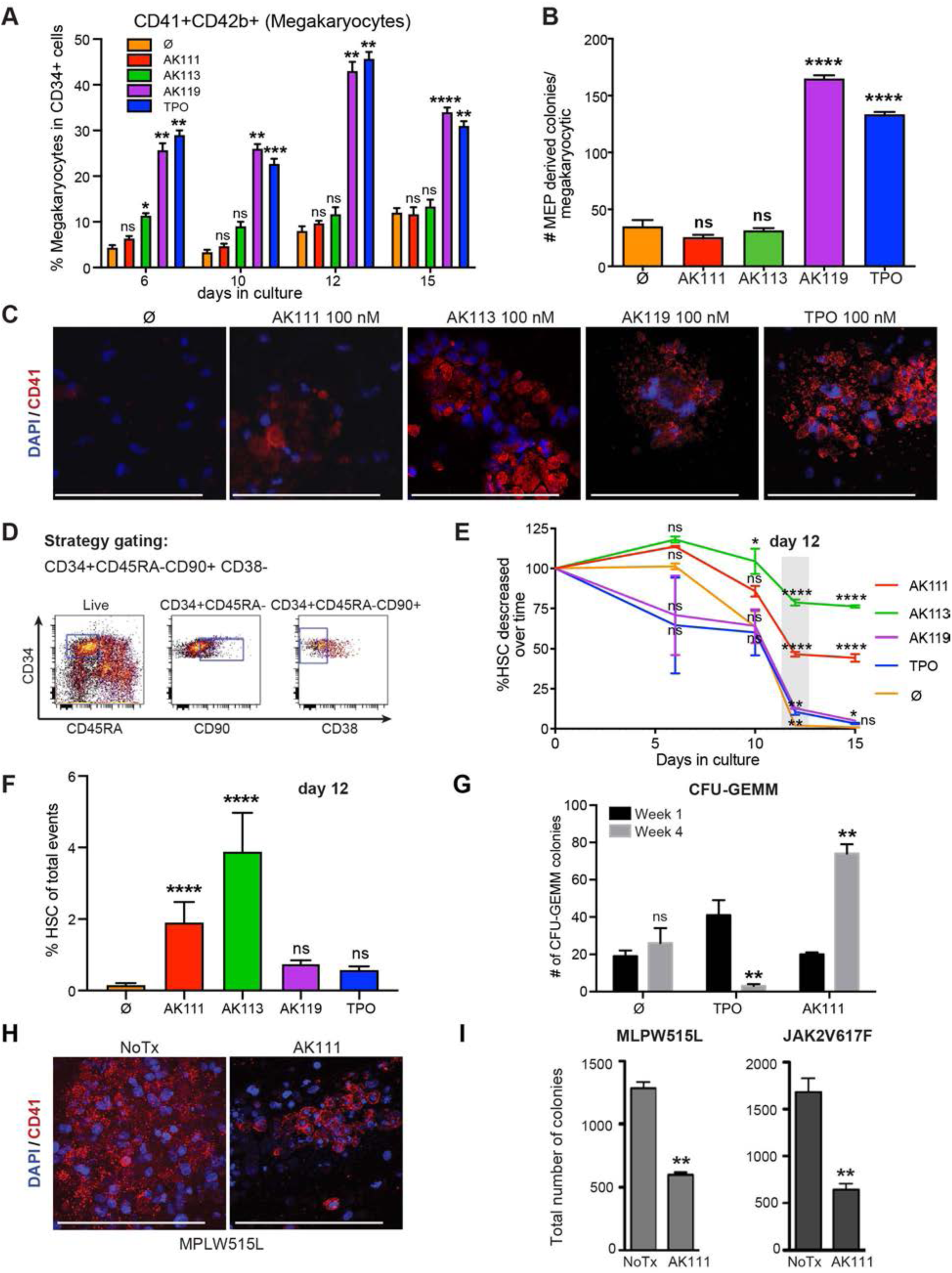
Diabodies split the dual function of TPO/TPO-R and block oncogenic signaling in Essential Thrombocytosis. (*A*) Percentage of megakaryocytes in CD34+ cells were quantified over 15 days in culture. Immune phenotypic characterization of liquid cultures of human HSC (HSC were sorted as Lin-CD34+CD45RA-CD90+CD38-cells on day 0) on indicated days plated under megakaryocytic differentiation inducing conditions with indicated cytokines or DBs (Ø: 20 ng/mL SCF was added to all conditions). Data are expressed as mean ± SD and analyzed by Ordinary two-way ANOVA, * *P* < 0.05, ** *P* < 0.01, *** *P* < 0.001, **** *P* < 0.0001, ns: non-significant. (*B*) Quantification of MEP-derived colonies in MegaCult cultures demonstrated minimal megakaryocytic differentiation with SCF and partial agonistic DBs AK111 and AK113, and full megakaryocytic differentiation with TPO and full agonistic DB AK119. Data are expressed as mean ± SD and analyzed by Ordinary one-way ANOVA, **** *P* < 0.0001, ns: non-significant. (*C*) Representative confocal images of MegaCult cultures treated with indicated DBs for 15 days, stained for megakaryocytic markers CD41 (red) and nuclear counter stain with DAPI (blue). Scale bar 100 µm. (*D*) Gating strategy for HSC isolation from human donor bone marrow (Live^+^/Lin^-^/CD34^+^/CD45RA^-^/CD90^+^/CD38^-^). (*E*) We measured the percent decrease of HSC at the indicated time points after treatment with Ø, AK111, AK113, AK119 or TPO over 15 days. We performed statistical analysis with One-way ANOVA followed by Dunnett’s multiple comparisons test with Ø as control group. * P < 0.05; ** P < 0.01; **** P < 0.0001, ns: non-significant. (*F*) Percent of HSC under Ø, AK111, AK113, AK119 and TPO induced culture on day 12. Data are expressed as mean ± SD and analyzed by One-way ANOVA, **** P < 0.0001, ns: non-significant. (*G*) HSC are harvested and treated for 12 days in liquid culture with the indicated cytokines or partial agonistic DB AK111 were subsequently placed in semisolid methylcellulose culture to evaluate their colony forming capacities over four rounds of weekly re-plating. Quantification of total numbers of CFU-GEMM colonies at the first and fourth re-plating for HSC previously treated with baseline SCF, TPO and AK111 in liquid cultures. Data are shown as mean ± SD. Unpaired t-test (** *P* < 0.01, ns: non-significant). (*H*) Representative confocal images of collagen-based cultures of MPLW515L mutated bone marrow of patients with essential thrombocythemia immune stained for CD41 (red) and DAPI (blue) (Scale bar 100 µm). (*I*) Quantification of collagen-based culture assays of MPLW515L, JAK2V617F, CALdel5, and CAL Ins52 mutated bone marrow of patients with essential thrombocythemia treated with indicated DB or without (NoTx). Data are shown as mean ± SD, statistical differences assessed by paired Student’s t-test (* *P* < 0.05. ** *P* < 0.01, *** *P* < 0.001, **** *P* < 0.0001).

Then, we evaluated the differential effects of DBs on HSC propagation and megakaryocytic differentiation when treated with DBs AK111 or 113 or 119 or TPO or low SCF (labeled ∅ control). We FACS purified single-cell HSC and MPP by their characteristic surface markers (HSC: lineage-CD34+CD38-CD45RA-CD90+, MPP: lineage-CD34+CD38-CD45RA-CD90-) (**Fig. 3*D, E***) at the indicated time points. In addition to the indicated factors, HSC were cultured over 15 days under conditions which strongly promoted megakaryocytic differentiation. We discovered that HSC cultured under megakaryocyte-inducing conditions but treated with AK113 or AK111 (partial TPO agonistic DBs) consistently preserved phenotypic HSC to a greater extent, 80% and ∼50% after 12 days in culture respectively, compared to only ∼10% in conditions treated with TPO or AK119 (full TPO agonistic DB), and very few in ∅ control (**Fig. 3*E* *and 3F***). Through colony forming and serial re-plating assays, we evaluated the ability of HSCs to self-renew and form multipotent colonies in vitro after treatment with TPO or DB AK111 and weekly cell re-plating over 4 rounds. Initially, we detected low numbers of CFU-GEMM colonies in AK113 and HSC treated with low concentrations of SCF without any additional growth factors or diabodies (∅), compared to 2-fold increased GEMM colony numbers in TPO-treated HSC after one round of re-plating. After 4 rounds of re-plating, however, HSCs treated with low SCF had low numbers of CFU-GEMM colonies whereas HSCs treated with TPO lost their GEMM colony forming capacity. In a stark contrast, HSCs treated with DB AK113 dramatically increased the number of CFU-GEMM colonies, demonstrating DB AK113s ability to enhance HSC stemness exhibited by increased serial re-plating in vitro (**Fig. 3*G***). In addition to effects on normal HSC we were curious whether partial agonistic DBs could interfere with oncogenic signaling in Essential Thrombocythemia (ET), a myeloid neoplasm and preleukemic condition with deregulated TPO signaling. Therefore we investigated whether antagonizing TPO signaling would have any therapeutic impact for ET. We assessed ex vivo colony formation from bone marrow samples of patients with ET carrying the MPL^W515L^ or the JAK2^V617F^ mutations. The bone marrow samples were grown without added cytokines or treated with the AK111 DB. We discovered that the AK111 DB not only significantly reduced colony formation in patients with MPLW515L and JAK2V617F mutated ET but also blocked differentiation towards megakaryocytes and platelets (**Fig. 3*H-I***).

### Partial Agonistic Diabodies propagate Hematopoietic Stem Cells In Vitro with Distinct Properties

To get a better understanding of the differential effects of diabodies on gene expression in HSC propagated in culture, we performed single-cell RNA sequencing of HSC treated with the DBs which we directly compared to native (=uncultured) HSC and MPP (**Fig. 4*A***). HSCs are defined as a quiescent stem cell population and maintain only low levels of the cell cycle gene TP53 (33). We confirmed extremely low expression levels of TP53 in native uncultured human HSC. We observed that HSCs-treated with DB AK111 or AK113 had TP53 levels even lower than native HSC stem cells. In contrast, HSC cultured solely with TPO or the full TPO agonistic DB AK119 had much higher expression levels of TP53 (**Fig. 4*A*)**. We also quantified proliferation by nuclear expression of MKI67 and found that HSC treated with AK111 and AK113 did not proliferate indicated by their low MKI67 which was similar to native HSC, while HSC treated with TPO or AK119 had a much higher MKI67 consistent with their active cell proliferation. Compared to native HSC, HSC cultured with AK111 and AK113 demonstrated similarly low MKI67 expression (**Fig. 4*A***). This was also the case for BAX, an apoptosis regulatory gene; native HSC and MPP were similar to AK111 and AK113 in their low BAX expression, which was somewhat higher in HSC propagated with TPO or AK119. UMAP analysis of single cell RNA seq data of cultured HSC revealed 10 subclusters amongst which three larger groups are discernable. Quite interestingly these data demonstrated that the full agonistic diabody AK119 clustered closely with TPO, while the partial agonistic diabody AK113 clustered in two distinct areas; one area next to TPO and SCF and another area far apart next to the more inhibitor diabody AK111. AK111 clustered in a spatially distinct area distant from TPO and AK119 (**Fig. 4*B*)**. These data support that the different diabodies induce distinct fate decisions in HSC.

**Fig. 4.**
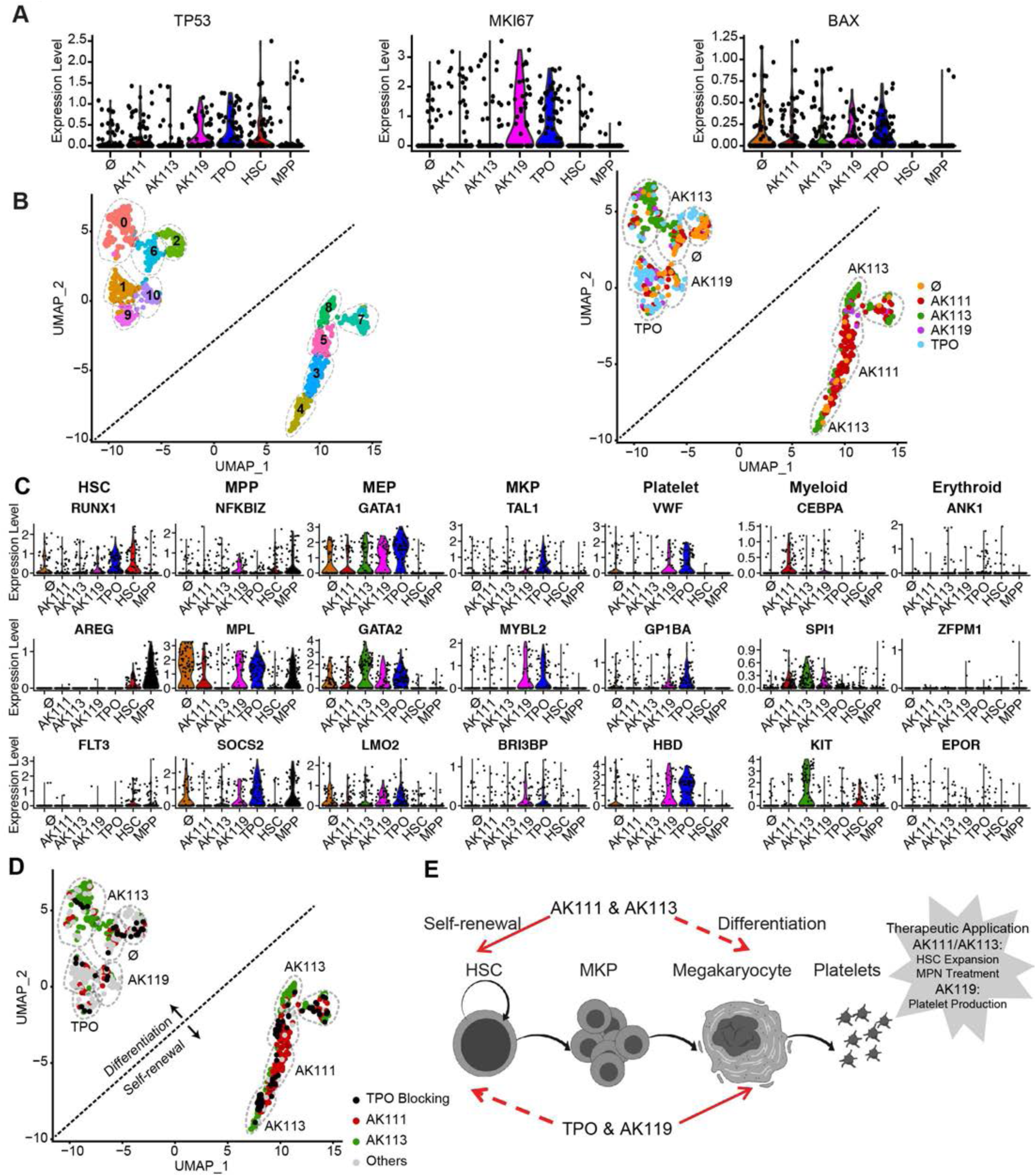
Partial agonistic diabodies propagate hematopoietic stem cells in vitro with distinct properties. (*A*) Single cell (sc) RNA-seq analysis shows the expression profiles of TP53, MKI67 and BAX (cell proliferation markers) in HSC after 12 days culture with Ø (20 ng/mL SCF) and combined with TPO DBs or TPO (100 nM). (*B*) Uniform manifold approximation and projection plots (UMAP) showing scRNA-seq data of FACS-isolated human HSC (CD34+CD45RA-CD90+CD38-) of 5 different culture conditions: Ø (20 ng/mL SCF) and combined with TPO DBs or TPO (100 nM). (*C*) Violin plots showing expression levels of genes critical for hematopoietic differentiation from HSC, MPP, MEP, MKP, platelet, mature myeloid and erythroid lineage stages for all diabody conditions on day 12 in culture. (*D*) Uniform manifold approximation and projection plots (UMAP) showing scRNA-seq data of FACS-isolated human HSC (CD34+CD45RA-CD90+CD38-) after treatment with blocking antibodies against TPO (*E*) Graphical schemata showing effects of diabodies on HSC proliferation and megakaryocytic differentiation.

Next we studied the change in gene expression profile along the differentiation path of HSCs to megakaryocytes in response to the treatment with different DBs compared to TPO (**Fig. 4C, S7, S8**). Quite comparable to TPO, AK119 activated transcriptional programs in primary HSC across the megakaryocyte differentiation spectrum including GATA1, GATA2 and LMO2, transcriptional programs characteristic for MEP stage, as well as programs pertinent for megakaryocyte precursors and platelets, TAL1, MYBL2 and VWF and HBD while inhibiting transcription factors driving myeloid and erythroid differentiation, CEBPA, KIT, ANK1 and EPOR. These data are in line with our findings described in Figure 1, which support that AK119 is a potent and full TPO agonist. Quite in contrast, AK111 which we showed to be the more inhibitory partial diabody agonist earlier in cell lines, appeared to dampen transcriptional activity throughout, only allowing for minimal expression of GATAs 1, 2 and selected myeloid transcriptional programs CEBPA and SPI1. Quite interestingly, AK113, the less inhibitory partial agonistic diabody in cell lines appeared to suppress MPL expression in primary cells the most and in a similar manner to that of native uncultured HSC, which is in line with data we showed earlier demonstrating that it was also the most potent in preserving the undifferentiated stage. In addition, AK113 suppressed megakaryocytic differentiation factors GATA 1 and 2 to a lesser degree while simultaneously allowing for expression of myeloid instructive programs KIT and SPI1 (**Fig. 4C, S7, S8**). One common feature of all TPO diabodies was the suppression of erythroid differentiation programs (e.g. ANK1, EPOR and ZFPM1 were not expressed), which was quite similar to that of native uncultured HSC and MPP. Thus, the partial agonist DBs show a surprising heterogeneity in induction of gene expression programs which could, in principle, lead to different functional effects.

### Partial Agonistic Diabodies Act by Reducing Signal Strength Through the TPO/TPO-R

We performed cell clustering of single-cell RNA-seq gene expression data and distinguished three populations labeled as 1, 2 and 3 (**Fig. 4*D*, S9**). Based on their single-cell RNA-seq expression profile, myeloid corresponded to cluster 1, HSC to cluster 2, and megakaryocytic and lineage+ to cluster 3. We evaluated the frequencies of these three cell populations treated with partial agonistic DBs in the presence or absence of a TPO-blocking antibody with single-cell RNA sequencing and compared the results to cells treated with SCF or TPO. HSC cultures treated with TPO contained a high percentage of cluster 3, lineage+ megakaryocytic cells, which decreased significantly with TPO blocking antibody. As expected, SCF mostly supported myeloid differentiation (cluster 1), while the cells treated with DB AK119 were quite similar to TPO supporting megakaryocytic differentiation (cluster 3). However, unlike TPO, a significant HSC proportion (cluster 2) was maintained. In general, HSCs treated with TPO-blocking antibody demonstrated a higher percentage of stem cells (cluster 2) and a lower percentage of megakaryocytic differentiated cells (cluster 3) compared to TPO (which showed the opposite). We observed a similar trend in HSC treated with DB AK111 and to a lesser extent with DB AK113, which retains a higher level of signaling activity (**Fig. S7**). UMAP plots in **Fig. 4*D*** demonstrate that treatment with DBs AK111 and with AK113 in a subset of cells appeared to have similar effects to the TPO-blocking antibody on HSC after 15 days of in vitro culture. Thus these data suggest that HSC maintenance/expansion by DBs AK111 and 113 is achieved by acting as a TPO dominant-negative by antagonizing TPO, as well as simultaneously delivering very weak agonistic signals. Collectively these data demonstrate that partially antagonizing TPO with DBs can fine tune the switch between HSC self-renewal and megakaryocytic differentiation and counteract oncogenic signaling in MPL and JAK2 mutated ET (**Fig. 4*E***).

## Discussion

Here we have explored the effects of manipulating TPO-R signaling in hematopoietic-fate decisions using surrogate TPO agonists and partial agonistic DBs. We have created a collection of TPO surrogate ligands with graded signaling potencies and strengths, enabling us to study this parameter on HSC biology. While all three DBs bound with comparable affinities to TPO-R, their downstream signaling outputs greatly differed and determined cell fate—maintenance of an HSC-like state with DBs AK111 and AK113 versus full maturation to megakaryocytes with AK119. While the structural basis of the differing signaling outputs elicited by the DBs is not resolved here, most likely the DBs bind to different epitopes on the TPO-R, resulting in a different range of dimeric TPO-R orientations and proximities. This is conceptually consistent with a previous study on DBs to EPO-R that elicited a similar pharmacological profile of full to partial agonism (28, 34). Structural analysis of the DB/EPO-R complexes revealed strikingly different EPO-R dimerization topologies that presumably impacted the orientation of the intracellular signaling machinery. Here we have extended this concept about basic mechanisms of cytokine-receptor signaling, to applications in HSC biology with therapeutic implications. This study, along with others demonstrating the effects of cytokine partial agonism on cytokine pleiotropy and function in systems such as IL-2, interferons, and stem cell factor, show protein engineering can generate partial agonists with unique pharmacology for dimeric receptors, borrowing conceptually from the GPCR structure-activity approaches, with distinctive functional properties that can be advantageous over wild-type (35–37).

Allogeneic HSC transplant is the only curative therapy for numerous hematologic malignancies and autoimmune diseases. While human HSC have been successfully purified since the 1980’s, their small cell numbers have limited their use in human transplant protocols. Purified HSC grafts would offer many advantages over CD34+-enriched stem-cell grafts. For example, purified HSC grafts do not confer acute and chronic Graft versus Host Disease (cGVHD), which is the leading cause of mortality and morbidity following a bone marrow transplant.

Despite substantial efforts to identify factors in the microenvironment of the bone marrow to support HSC stemness (3, 4, 6, 7), it has not as yet been possible to stably expand purified human HSC ex vivo for clinical use despite preclinical work demonstrating that HSC can be propagated in culture with pyrimidoindole-derived molecules, HOXB or by suppressing the aryl hydrocarbon receptor (38–41). In this study we demonstrated that partial agonistic DBs AK111 and AK113 allow for HSC maintenance and expansion ex vivo by diminishing and/or modulating activating downstream signals through TPO-R. In contrast to these other approaches which act through obscure mechanisms, TPO DBs modulate signaling solely through the TPO-R, thus side-effects are anticipated to be minimal. TPO-DBs are able to deliver customized intensities of downstream TPO-R signals, and thus mimic the physiological conditions of the TPO regulation while more tightly controlling downstream signaling output. In addition, the TPO DBs split the functions of TPO with regard to megakaryopoiesis and HSC expansion, a unique feature, which is not a property of any of the available TPO mimetics nor the above-mentioned stem-cell expanding experimental reagents. The dual function of TPO is an unanswered and burning open question in the otherwise well-studied field of TPO signaling. Thus, TPO DBs also represent an interesting reagent for further exploratory research in this area.

The importance of cytokines in maintaining the hematopoietic stem-cell pool and cell lineage specifications has been well established in the literature. For instance, SCF and TPO are required cytokines for HSC. While HSCs generated by DBs show strong gene expression similarities with uncultured HSCs by single-cell RNA seq, they also have very specific features. HSC_AK111_ expressed increased IL6R and decreased FLT3 ligand compared to uncultured HSC. It is understood that the expression of FLT3 increases between the HSC and MPP stage and plays a key role for multipotent and lymphoid progenitors and myeloid cells. While the absence of FLT3L does not alter megakaryocyte-erythroid progenitor development, the cytokine TPO has been shown to potentiate FLT3-mediated effects. Our single-cell RNA seq data show that HSCs cultured with partial inhibitory TPO DBs and very low SCF lacked FLT3 expression and retained at least short-term stem-cell-like properties; thus, these culture conditions appear optimal for stem-cell maintenance.

Using single-cell transcriptomics (scRNA-seq) in conjunction with protein expression studies, we confirmed that the stem cells generated by DBs (which we found partially block the activity of MPL) produce an original pool of stem cells in vitro (38, 39). By clustering HSC and MPP we can conclude that after nearly two weeks of culture, the HSC_AK111_ populations are closer to HSC than MPP populations. For example, the HSC_AK111_ demonstrate a TP53 expression pattern very similar to that of uncultured HSC. We know that TP53 expression is tightly controlled in HSCs, and it has been shown that the Trp53, p16, p19 triple mutant allows long-term hematopoietic reconstitution (40), and HSC expansion (41). Our studies show a basal low level of TP53 in HSC AK111 quite similar to HSC.

More recently, it has been demonstrated that mutations in the cytokine TPO reduce engraftment capacity, further demonstrating the essential role of TPO for primary hematopoiesis (42, 43). The use of TPO as a drug initially had been quite limited due to antibody formation and thrombocytopenia. Nevertheless, the second generation of TPO receptor agonists are now being used successfully in the clinic for severe immune thrombocytopenia and bone marrow failure syndromes and are under evaluation for a wide range of other thrombocytopenic disorders, including those associated with chemotherapy and myelodysplastic syndromes. Thus, while generally much better tolerated, unwanted side effects still occur with second generation TPO mimetics including thrombocytosis, thrombosis, bone marrow reticulin fibrosis, rebound thrombocytopenia and liver toxicity. DBs deliver different intensities of downstream MPL signals, and thus mimic the physiological conditions of TPO regulation while more tightly controlling downstream signaling output. In addition to the above discussed applications, we found this to be particularly relevant to oncogenic signaling downstream of the TPO-R as they occur in ET, a myeloproliferative neoplasm. To our great surprise we discovered that partial agonistic TPO DB blocked oncogenic colony formation in patients with ET by decreasing downstream signaling of TPO-R. Perceivable possible clinical applications of TPO DB could be conditions requiring full or partial TPO agonism such as platelet production, HSC expansion/ maintenance, as well as antagonizing oncogenic signaling in ET.

## Acknowledgements

K.C.G. acknowledges NIH-RO1-AI51321, Ludwig Institute, Mathers Foundation, and HHMI for funding: G.W.: NHLBI, Ludwig Institute, SRF and Goldman-Sachs Foundations, DFG for salary support of TL.

## Authors contributions

Conceptualization, G.W. and K.C.G; Methodology, G.W., I.M. Investigation, G.W., I.M., L.C., T.L., C.V.N., S.W., N.T., M.G., S.D., M.E., N.T. Writing – Original Draft, G.W. Writing – Review & Editing, G.W., K.C.G., I.H., J.P., I.M., L.C. Funding Acquisition, K.C.G., G.W. Resources, J.G., S.Q., C.C. Supervision, K.C.G., G.W.

## Declaration of Interest

The authors declare no competing interests.

## Supplementary Information

### SI Materials and Methods

#### TPO Diabodies Sequence

pAK111 - patent: US2012/0053326 – **heavy** and light chain clone 6-4-50> (pg 22) MLLVNQSHQGFNKEHTSKMVSAIVLYVLLAAAAHSAFAGS

**EVQLVESGGGLVQPGRSLRLSCATSGFTFDNYAMYWVRQAPGKGLEWVSGISWNSGDIGYADSVKGRFTISRDNAKNSLYLQMNSLRAEDTALYYCARDAGFGEFHYGLDV WGQGTTVTVSS**GGGGSAIQLTQSPSSLSASVGDRVTITCRASQGISSALAWYQQKPG KVPKLLIYDASSLESGVPSRFSGSGSGTDFTLTISSLQPEDFATYYCQQFNSYPWTFG QGTKVEIKR AAAHHHHHHHH

pAK113-patent: US2012/0053326 – **heavy** and light chain clone 7-10> (pg 20) MLLVNQSHQGFNKEHTSKMVSAIVLYVLLAAAAHSAFAGS

**EVQLVESGGGLVQPGRSLRLSCAASGFTFDDYAMHWVRQAPGKGLEWVSGISWNSGSIGYADSVKGRFTISRDNAKNSLYLQMNSLRAEDTALYYCAKNLWFGEFRYWYFDL WGRGTLVTVSS**GGGGSAIQLTQSPSSLSASVGDRVTITCRASQGISSALAWYQQKPG KAPKLLIYDASSLESGVPSRFSGSGSGTDFTLTISSLQPEDFATYYCQQFNSYPLTFGG GTKVEIK AAAHHHHHHHH

pAK119: **heavy** and light chain (26)

MLLVNQSHQGFNKEHTSKMVSAIVLYVLLAAAAHSAFAGS

**QVQLQQSGPELVKPGASVKISCKASGYAFSSSWMNWMKQRPGKGLEWIGRIYPGD GDTNYNGKFKGKATLTADKSSSTAYMQLSSLTSEDSAVYFCARARKTSWFAYWGQ GTLVTVSA**GGGGSDIVLTQSQKFMSTSVGDRVSISCKASQNVGNIIAWYQQKPGQSP KALIYLASYRYSGVPDRFTGSGSGTD<u>FTLTISNVQSED</u>LAEYFCQQYSSSPLTFGAGTK LEIK AAAHHHHHHHH

#### Protein Expression and Purification

TPO diabodies were manufactured as recently described (23, 26), cloned into the pAcGP67-A vector (BD Biosciences) in frame with an N-terminal gp67 signal sequence and a C-terminal hexahistidine tag and produced using the baculovirus expression system, as described by LaPorte et al.(44). Baculovirus stocks were prepared by transfection and amplification in Spodoptera frugiperda (Sf9) cells grown in SF900II media (Invitrogen) and protein expression was carried out in suspension Trichoplusiani (High Five) cells grown in InsectXpress media (Lonza). Following expression, proteins were captured from High Five supernatants after 60 hr by nickel-NTA agarose (QIAGEN) affinity chromatography, concentrated, and purified by size exclusion chromatography on a Superdex 200 column (GE Healthcare), equilibrated in 10mMHEPES (pH 7.2) containing 150mMNaCl and 15% glycerol. Recombinant diabodies were purified to greater than 98% homogeneity.

#### Phospho-Flow Cytometry

Intracellular staining of pSTAT5 was performed after permeabilization of UT7 TPO-R cells with ice-cold methanol (100%, v/v). Alexa Fluor 647–conjugated antibodies specific for pSTAT5 were purchased from Cell Signaling (CST) and used at a 1:50 dilution. The extent of STAT5 phosphorylation was calculated by subtracting the mean fluorescence intensity (MFI) of pSTAT5 of the stimulated samples from that of the unstimulated sample. The normalized values were plotted against cytokine concentration to yield dose-response curves from which the EC_50_ values were calculated on the basis of nonlinear, least squares regression fit to a sigmoidal curve using graphpad prism software.

#### Cell Proliferation

Two thousand UT7 TPO-R cells per well were seeded in a 96-well plate and stimulated with the indicated concentrations of TPO or Diabodies. After 96 hours of stimulation, the cells were harvested, and cell number was determined by luminescence using a Promega cell counting kit. The number of cells obtained for each agonist was plotted against the cytokine concentration to obtain sigmoidal dose-response curves, from which the EC_50_ values for UT-7 hTPO-R cell proliferation was calculated.

#### TPO-R Diabodies Binding Experiments

The Saccharomyces cerevisiae strain EBY100 was transformed with the pCT302_TPO-R SD1-2 vector and grown for 2 days at 30°C on selective dextrose casamino acids (SDCAA) media, followed by induction in selective galactose casamino acids (SGCAA) medium (pH 4.5) for 2 days at 20°C. Yeast were then incubated with the indicated concentrations of biotinylated TPO-R Diabodies for 1 hour at 4C, followed by Streptavidin-Alexa647 (1:200 dilution) for 15 min at 4C. Fluorescence was analyzed on an Accuri C6 flow cytometer.

#### Primity Bio Pathway Phenotyping

HSCs and MPP cells were starved overnight, then stimulated with saturated concentrations of TPO and indicated diabodies for 15, 60, and 120 min, and then fixed with 1% PFA for 10 min at room temperature. The fixed cells were prepared for antibody staining according to standard protocols. After, the fixed cells were permeabilized in 90% methanol for 15 min. The cells were stained with a panel of antibodies specific to the markers indicated (Primity Bio Pathway Phenotyping service) and then analyzed on an LSRII flow cytometer (Becton Dickinson). The Log2 Ratio of the median fluorescence intensities (MFI) of the stimulated samples were divided by the unstimulated control samples to be calculated as a measure of response.

#### Single-Molecule Fluorescence Imaging

Single-molecule imaging experiments were conducted by total internal reflection fluorescence (TIRF) microscopy with an inverted microscope (Olympus IX71) equipped with a triple-line total internal reflection (TIR) illumination condenser (Olympus) and a back-illuminated electron multiplied (EM) CCD camera (iXon DU897D, Andor Technology) as described in more detail previously (28, 45). A 150 × magnification objective with a numerical aperture of 1.45 (UAPO 150× /1.45 TIRFM, Olympus) was used for TIR illumination of the sample. All experiments were carried out at room temperature in medium without phenol red, supplemented with an oxygen scavenger and a redox-active photo protectant to minimize photobleaching (46). For cell surface labeling of mXFP-tagged receptors, DY647- and Rho11-labeled NBs were added to the medium at equal concentrations (2 nM each). The labeled NBs were kept in the bulk solution during the whole experiment in order to ensure high equilibrium binding. TPO and diabodies were incubated for at least 5 min prior to image acquisition. Rho11 and TMR were excited by a 561 nm laser (CrystaLaser) at 0.95 mW (∼32 W/cm^2^) and DY647 by a 642 nm laser (Omicron) at 0.65 mW (∼22 W/cm^2^). Fluorescence was detected using a spectral image splitter (DualView, Optical Insight) with a 640 DCXR dichroic beam splitter (Chroma) combined with the bandpass filter 585/40 (Semrock) for detection of Rho11 and 690/70 (Chroma) for detection of DY647 dividing each emission channel into 512 x 256 pixels. Image stacks of 150 frames were recorded at a time resolution of 32 ms/frame for each cell.

#### Single-Molecule Analyses

Single-molecule localization and tracking was carried out using the multiple-target tracing (MTT) algorithm (47) as described previously (48). Immobile molecules were identified by spatiotemporal cluster analysis (49) and removed from the dataset (typically ∼15-20% of all localizations) prior to quantifying diffusion and dimerization. Receptor dimerization was quantified based on sequential co-localization and co-tracking analysis as described in detail recently (45). After aligning Rho11 and DY647 channels with sub-pixel precision by using a spatial transformation based on a calibration measurement with multicolor fluorescent beads (TetraSpeck microspheres 0.1 µm, Invitrogen), individual molecules detected in both spectral channels of the same frame within a distance threshold of 100 nm were considered co-localized. For single-molecule co-tracking analysis, the MTT algorithm was applied to this dataset of co-localized molecules to reconstruct co-locomotion trajectories (co-trajectories) from the identified population of co-localizations. For the co-tracking analysis, only co-trajectories with a minimum of 10 consecutive steps (∼300 ms) were considered. This cut-off was determined based on systematic analysis of a negative control experiment with non-interacting model transmembrane proteins (45) in order to minimize background from random co-localization.

#### Ba/F3-MPL, UT7-MPL Cell Line Generation and Immunoprecipitation

Wild-type human MPL (NM_005373.1) was cloned into the retroviral expression vector pMX-puro using EcoRI and XhoI cloning sites. The MPL intracellular domain point mutations (Y591F, Y625F and Y630F) were introduced using the QuikChange mutagenesis kit (Stratagene). Ba/F3-MPL cells lines were generated by retroviral transduction as described previously (32). Prior to stimulation, Ba/F3-MPL cells were cytokine starved for 16hrs. After Diabodies or TPO stimulation, cell lines were lysed in Nonidet P40 (NP-40) lysis buffer (50mM tris(hydroxymethyl)aminomethane-HCl, pH 7.4, 1% NP-40, 150mM NaCl, 1mM ethylenediaminetetraacetic acid, 10mM β-glycerolphosphate, 1mM Na3VO4, 10mM NaF) containing 1% protease inhibitors (Sigma-Aldrich). Supernatants were precleared by preincubating for 1 hour at 4°C with protein A beads (Millipore) before MPL antibody (rabbit anti-MPL/TPO-R; Millipore) incubation overnight. Immunoprecipitates were washed before resuspension in Bolt™ LDS Sample Buffer (Thermo Fisher) and heated for 10 minutes at 70°C. Immune precipitates were fractionated by sodium dodecyl sulfate (SDS)–polyacrylamide gel electrophoresis and transferred to polyvinylidene fluoride membranes. Protein expression was detected by incubating with anti-phospho-tyrosine antibody (4G10-platinum, mouse anti-phospho-tyrosine; Millipore) and visualized by chemiluminescent detection reagent (ECL-plus; GE Healthcare) and quantified by densitometry using ImageJ analysis software (National Institutes of Health, http://rsbweb.nih.gov/ij).

#### Isolation and Culture of HSC from CD34 Enriched Human Bone Marrow

Growth protocol. Human bone marrows were obtained from Stanford Medical Center with informed consent, according to Institutional Review Board (IRB)-approved protocols (Stanford IRB 39881). CD34+ progenitors from each sample were isolated by Ficoll separation and CD34+ progenitors were isolated by a positive selection using an immunomagnetic cell sorting system (AutoMacs; Miltenyi Biotec, Bergisch Gladbach, Germany). 56,000 cells were incubated in IMDM media supplemented with 10% BSA, 1% L-glutamine, 450 µM MTG, 1% ITG, liposomes, 1% penicillin/streptomycin, in the presence SCF 20 ng/ml and TPO or the different diabodies at 100 nM, and/or blocking antibody against TPO (R&D AF-288-NA) for 12 days, with complete change of medium every other day. After 12 days, cells were pelleted and resuspended in PBS 2% FCS. Fc block was performed for 20 min and the mononuclear cells were stained with directly conjugated anti-mouse CD34, CD38, CD45RA, CD90 and HSC were isolated as DAPI-CD34+CD38-CD45RA-CD90+ cells on a BD FACS ARIA sorter.

#### Single Cell RNA Sequencing

Single-cells were collected in lysis buffer in 96-well plates, followed by reverse transcription with template-switch using an LNA-modified template switch oligo to generate cDNA. After 21 cycles of pre-amplification, DNA was purified and analyzed on an automated Fragment Analyzer (Advanced Analytical). Each cells cDNA fragment profile was individually inspected and only wells with successful amplification products (concentration higher than 0.06 ng/ul) and with no detectable RNA degradation were selected for final library preparation. Fragmentation assays and barcoded sequencing libraries were prepared using Nextera XT kit (FC-131-1024; Illumina) according to the manufacturer’s instructions. Barcoded libraries were pooled and subjected to 75 bp paired-end sequencing on the Illumina NextSeq instrument. The single cell RNA sequencing and data analysis was performed and analyzed as previously described by the Quake laboratory (50).

#### Cell Culture with Diabodies

CD34+ Bone Marrow derived cells were cultured in suspension in IMDM media supplemented with 10% BSA, 1% L-glutamine, 450 µM MTG, 1% ITG, liposomes, 1% penicillin/streptomycin and baseline SCF at 20 ng/ml for 12 days. Additional conditions received 100 nM of TPO, or 100 nM of different Diabodies agonists like AK111 and AK113, or antagonist AK119, with or without 14 µg/ml of TPO blocking antibody. Media was changed every other day completely.

#### Serial Re-plating Assay

Myeloid colony–forming assays were performed in methylcellulose-based medium (M3232) as per manufacturer’s protocols (Stem Cell Technologies, Vancouver, British Columbia, Canada). Cells were plated at 1.5×10^4^ cells per dish in duplicate, with media plus 20 ng/mL SCF, or TPO or Diabody AK111 at 100nM. Colony forming unit-granulocyte, erythrocyte, macrophage, megakaryocyte (CFU-GEMM) were counted at day 7, re-passaged weekly until week 4 in Methocult at 1.5×10^4^ cells.

#### Accession Numbers

RNA-seq data can be accessed via Gene Expression Omnibus (GEO) at National Center for Biotechnology Information (NCBI) under the accession number GSE115235.

#### Statistics

Statistical analyses were performed using Prism software (GraphPad Software). Statistical significance was determined by the unpaired Student’s t test for comparisons between two groups and one-way ANOVA for multigroup comparisons (n.s. non-significant; *P* > 0.05; * *P* < 0.05; ** *P* < 0.01; *** *P* < 0.001; **** *P* < 0.0001). In statistical graphs, points indicate individual samples, and results represent the mean ± SD unless indicated otherwise.

#### Study Approval

De-identified patient specimens in paraffin and discarded fresh patient tissues were used for our studies as approved in IRB-39881. Mice were maintained in Stanford University Laboratory Animal Facility in accordance with Stanford Animal Care and Use Committee and National Institutes of Health guidelines (SU-APLAC 30911 and 30912).

## SI Figures

**Figure S1.**
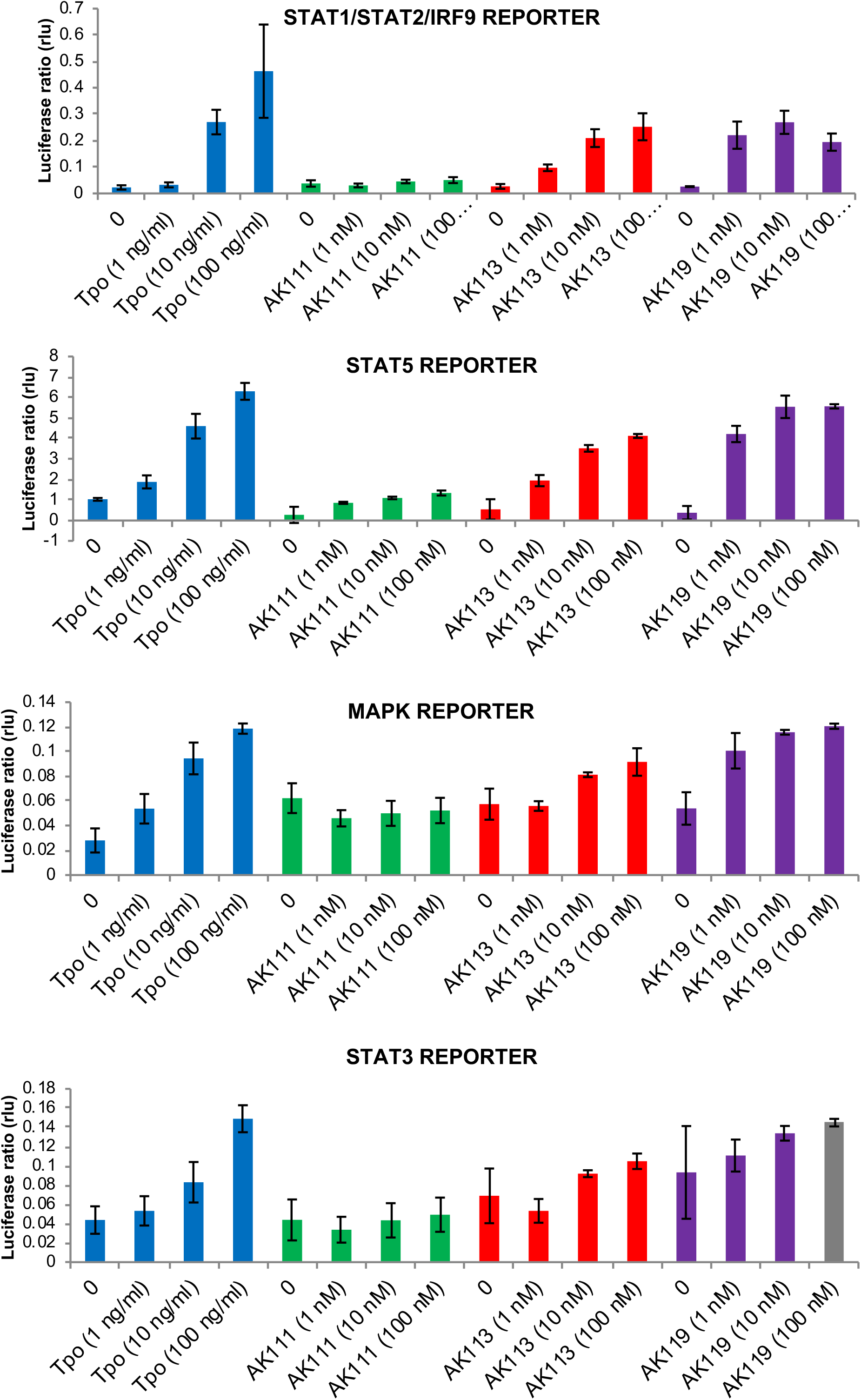
Reporter assays for STAT1/STAT2/IRF9, STAT5, MAPK and STAT3 demonstrated that AK111 blocked signaling through STAT1/STAT2/IRF9, and STAT5, and reduces signaling activity through STAT3, while no changes have been noticed for the MAPK reporter activity. Of note, AK113 demonstrated somewhat attenuated response to STAT signaling, and AK119 showed significant increased signaling activities through these pathways albeit at much lower concentrations. Data are shown as mean ± SD.

**Figure S2.**
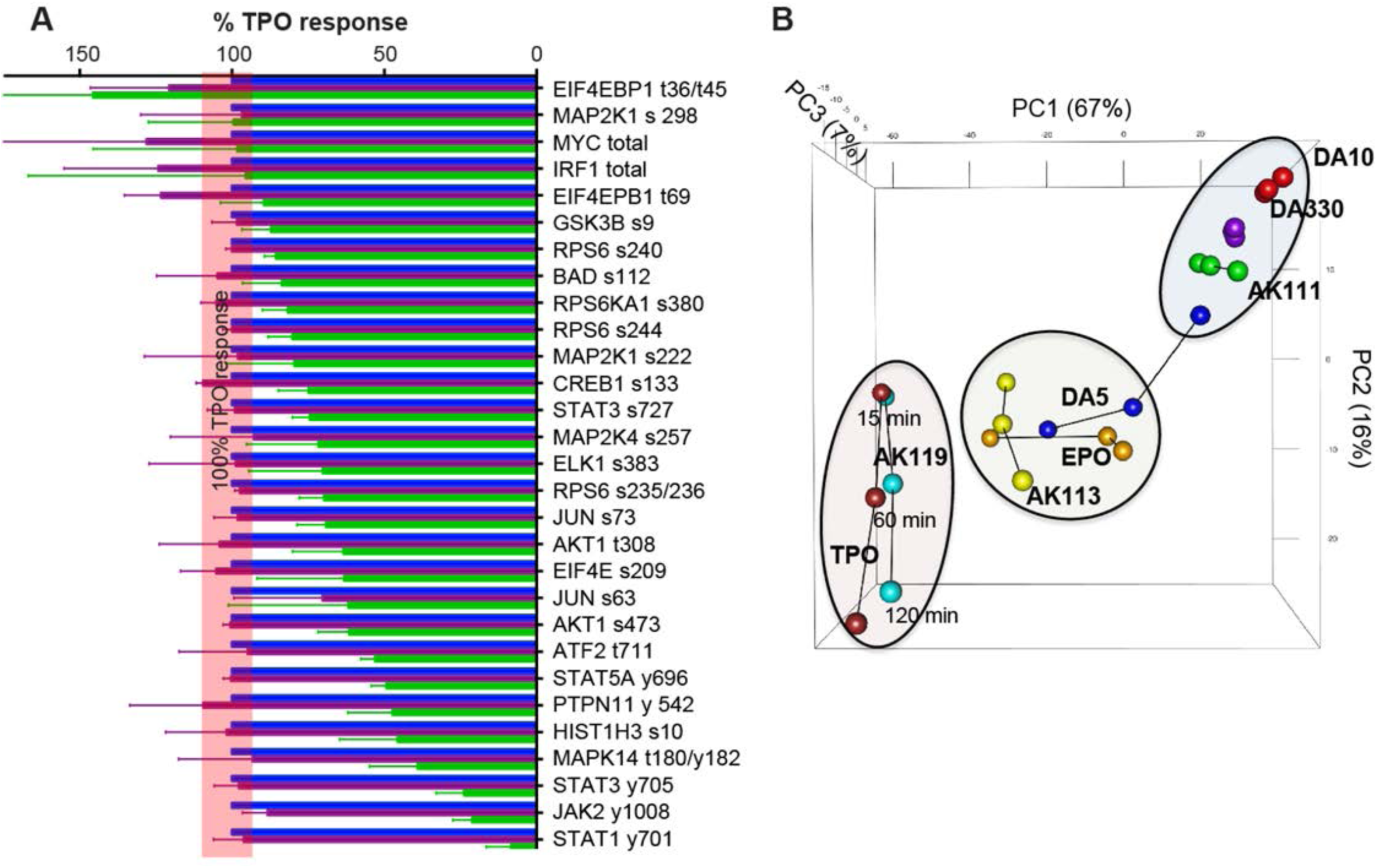
(*A*) The levels of signal activation induced by TPO and partial and full agonistic DBs at 15 min of stimulation were normalized to those induced by TPO and aligned based on signaling potency. The red line represents the TPO-signaling activation potency normalized to 100%. Data mean ± SD are from three independent replicates. (B) PCA of signaling data directly compared TPO and TPO DBs (AK111, AK113 and AK119) at 15 min, 60 min and 120 min of stimulation to previously published EPO and EPO DBs (DA5, DA10 and DA330) data.

**Figure S3.**
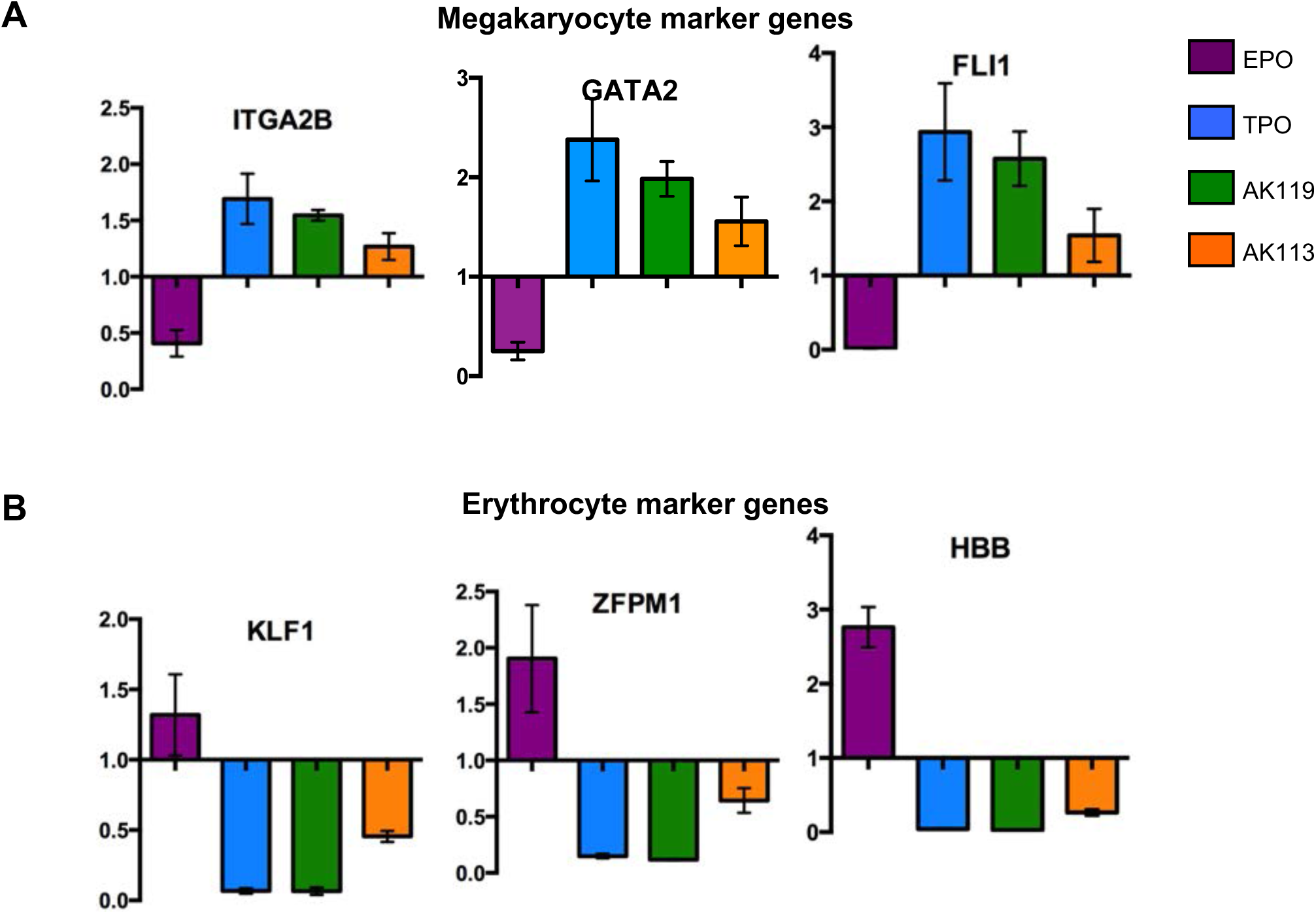
Gene expression profiles of indicated megakaryocytic lineage (A) and erythroid lineage (B) defining transcription factors profiled by qPCR in UT-7 hTPO-R expressing cells after treatment with EPO, TPO or the indicated diabodies for 21 days. Data are shown as mean ± SD.

**Figure S4.**
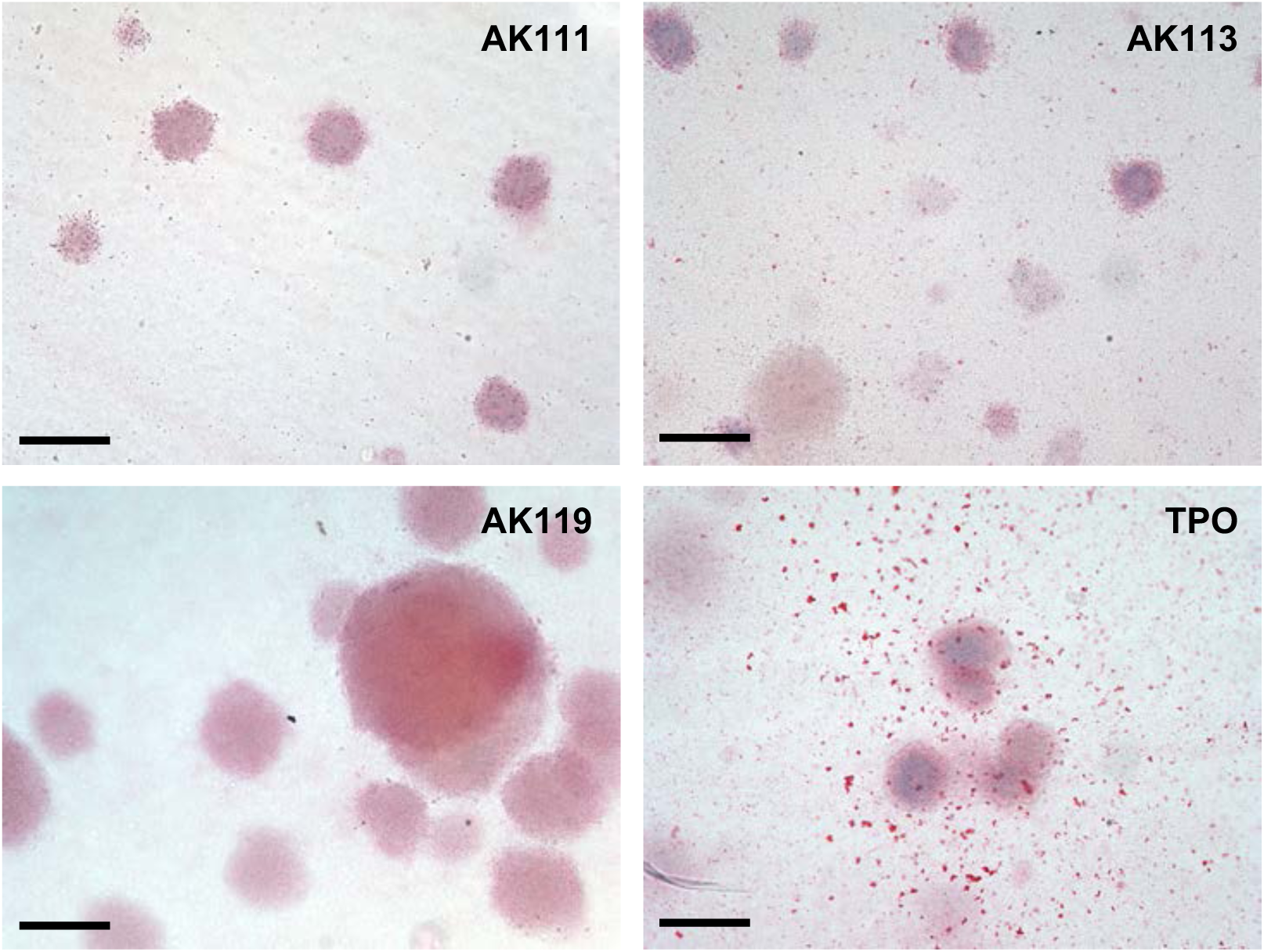
MEP cells were FACS purified from human bone marrow, and subsequently seeded in Megacult collagen-based media containing the indicated diabodies or TPO to test differentiation potential of diabodies of MEP progenitors. The number and identity of the colonies were assessed at day 15 by an experienced double blinded hematopathologist. In addition, cultures were also stained for CD42b, a lineage marker for megakaryocytic differentiation. These assays were performed in triplicate and repeated once. Scale bar: 100 μm.

**Figure S5.**
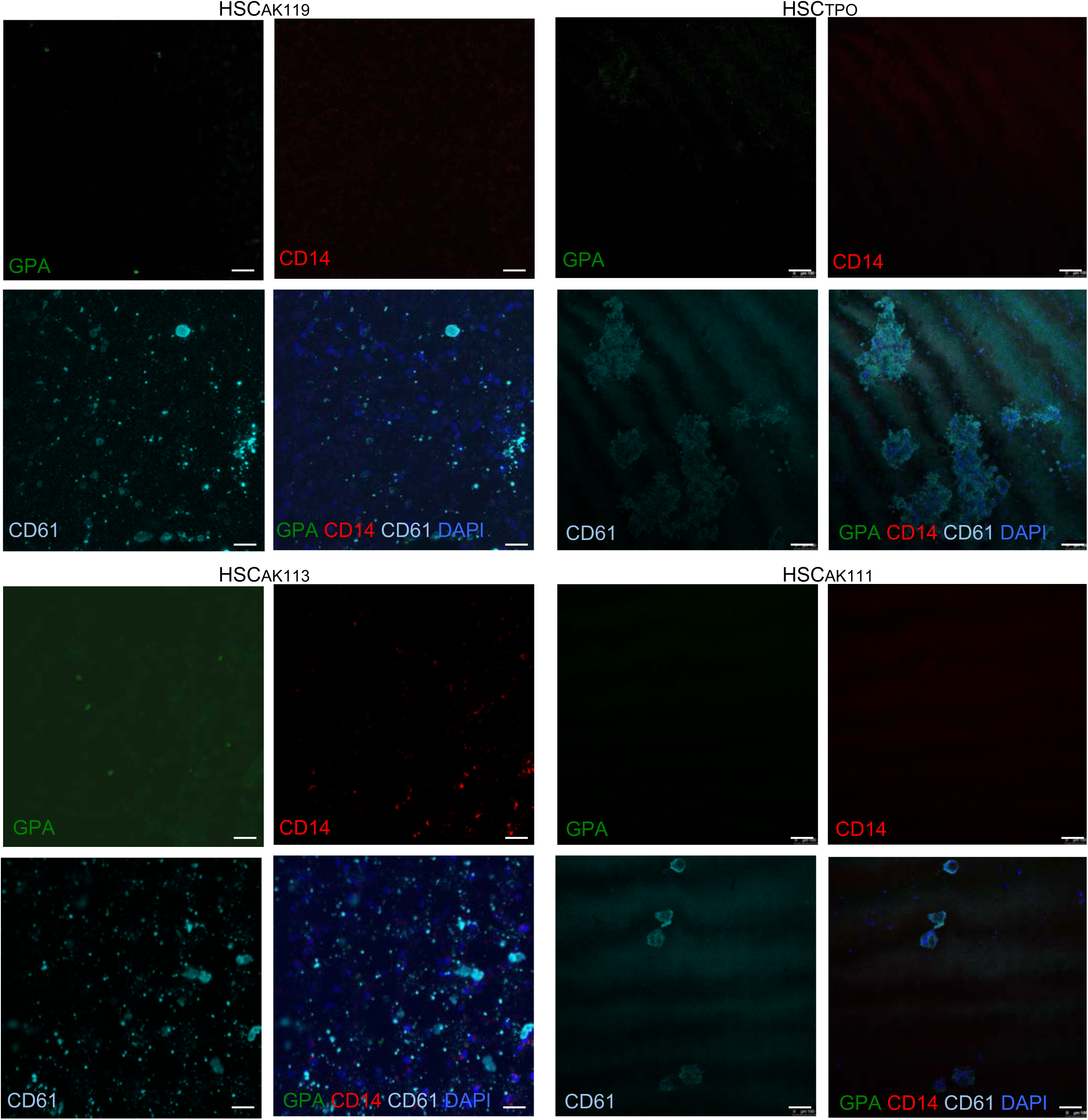
Confocal images of FACS purified and seeded HSC undergoing megakaryocytic differentiation in Mega-Cult supplemented with the indicated diabodies for 15 days. While AK119 fully promoted differentiation to megakaryocytic cells similarly to TPO, as indicated by polyploid large cells with positive expression of CD41 and/or CD61, AK111 and AK113 significantly prevented it (no significant larger polynucleated megakaryocytic cells, instead predominantly high N/C ratio mononuclear cells expressing CD61. Of note, no significant contribution to erythroid lineage (assessed by GPA staining), nor the myelomonocytic lineage (assessed by CD14 staining) was observed in semisolid cultures. Scale bar: 100 μm.

**Figure S6.**
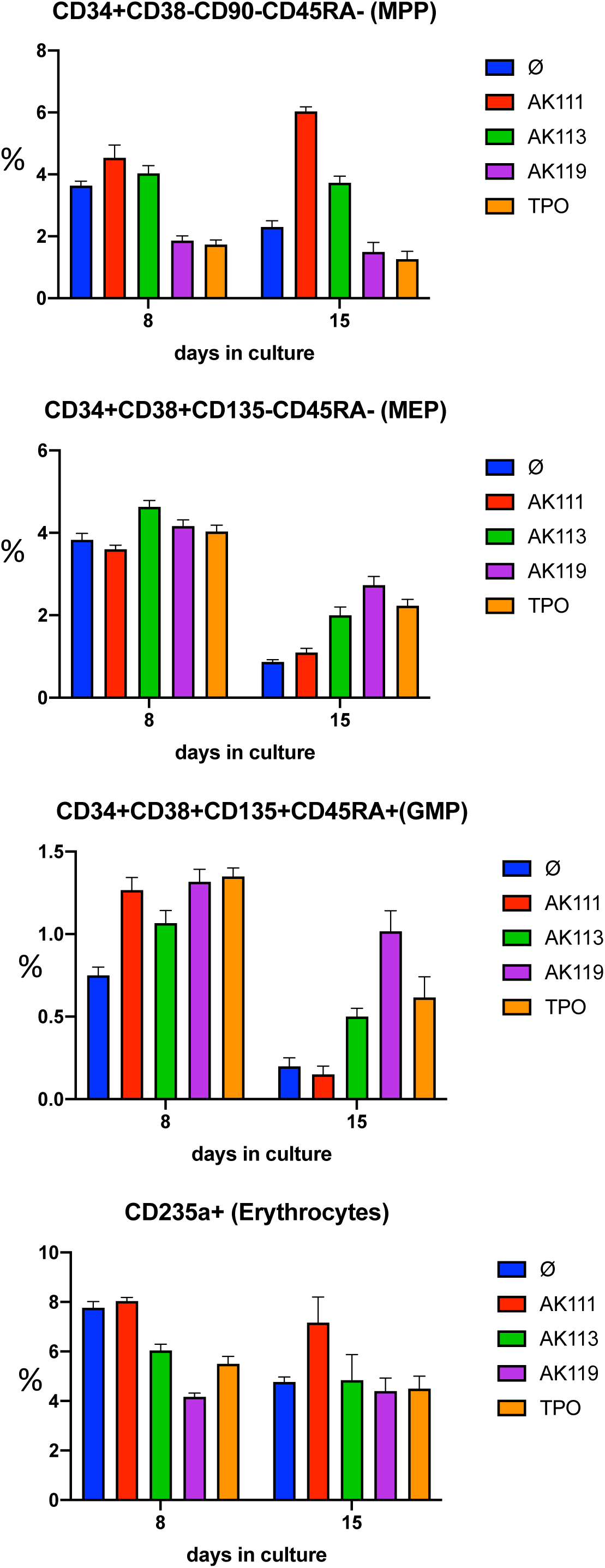
Flow cytometry analysis of liquid cultures of sorted human HSC treated with 20 ng/mL SCF only as culture baseline (Ø), the indicated diabodies or TPO over 15 days. No significant contribution to myeloid progenitors (MPP), Megakaryocytic-Erythroid progenitors (MEP), Granulocyte-Monocyte progenitors (GMP), or the erythroid lineage was appreciated in liquid culture under megakaryocytic differentiation conditions over 15 days in culture. The data are shown as relative percentage per total cell events on day 8 and 15.

**Figure S7.**
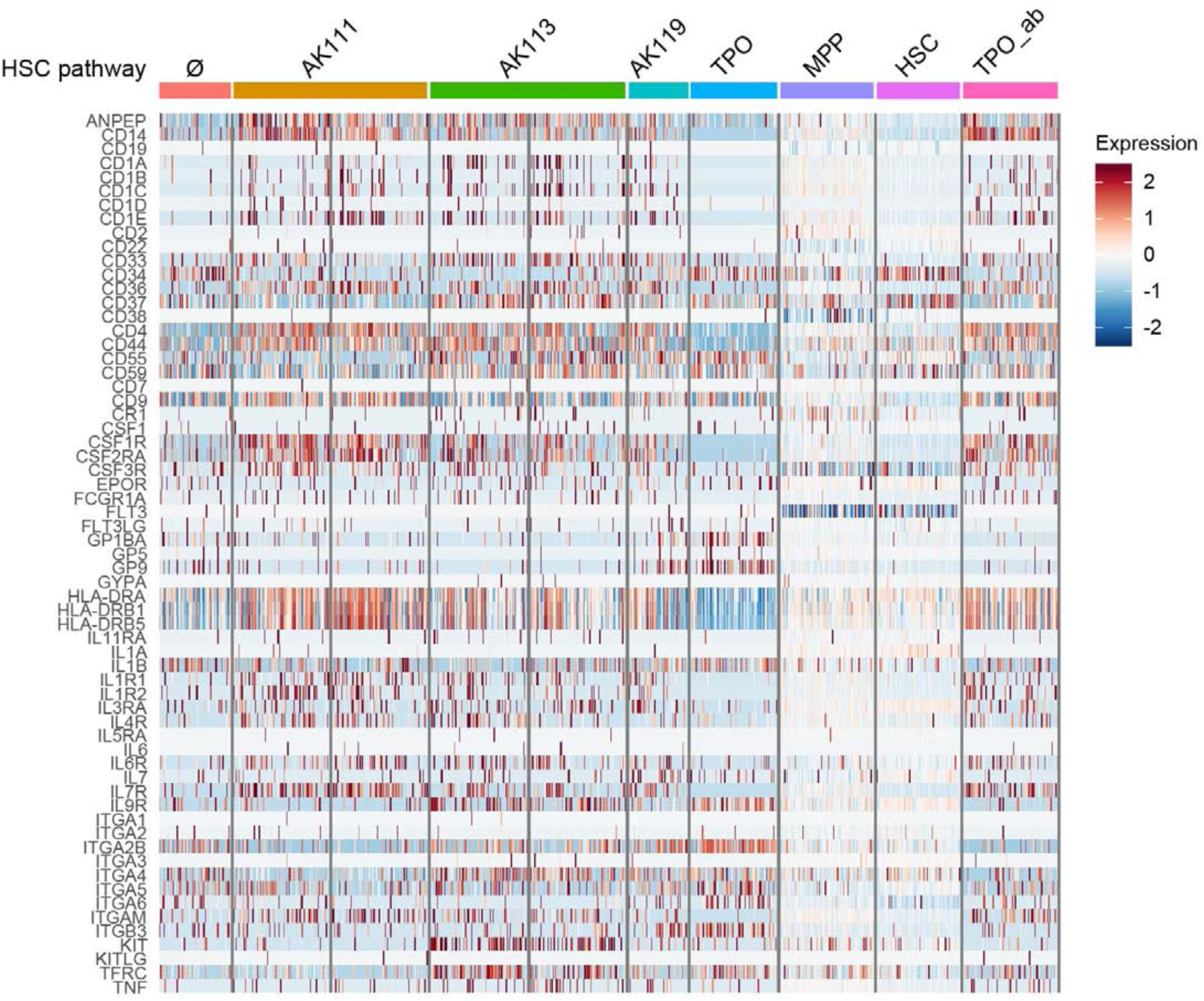
Heatmaps demonstrating unbiased clustering of single cell RNAseq of early hematopoietic genes differentially regulated by diabodies.

**Figure S8.**
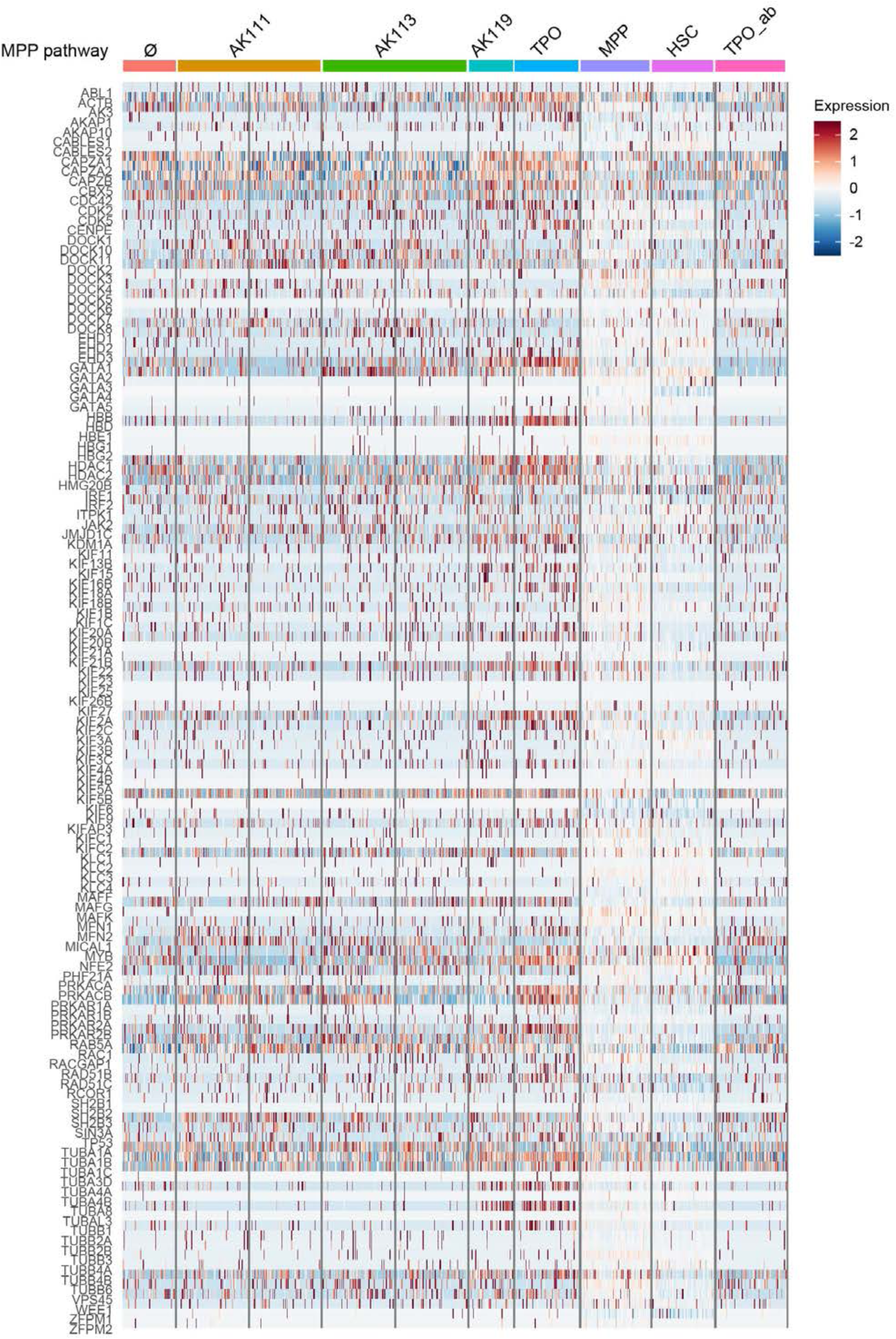
Heatmaps demonstrating unbiased clustering of single cell RNAseq of precursor genes differentially regulated by diabodies.

**Figure S9.**
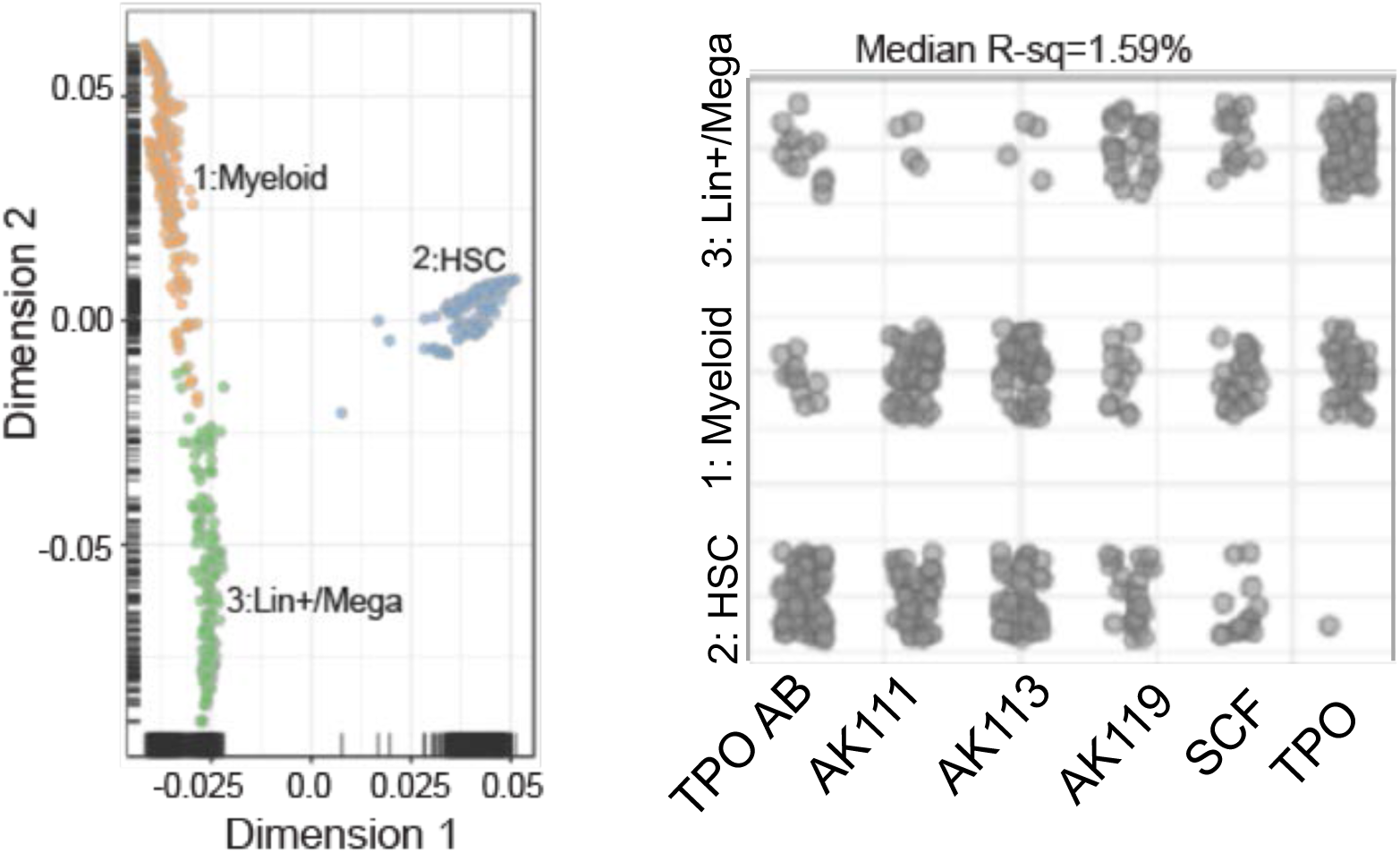
(A) Spatial resolution of single cell RNA sequencing data of human HSC (Lin-CD34+CD45RA-CD90+CD38-) treated with TPO blocking antibody (TPO_AB), the indicated diabodies −/+ blocking TPO antibody, baseline SCF (20 ng/ml), TPO (100 nM) −/+ blocking TPO antibody distinguished three main clusters, cluster 1 - myeloid lineage, cluster 2 - HSC, and cluster 3 - lineage+ megakaryocytic (Lin+Mega). (B) Individual clusters of sorted HSC (Lin-CD34+CD45RA-CD90+CD38-) after cultured with indicated diabodies for 12 days with and without TPO-blocking antibody (TPO_AB).

## Notes

### Competing Interest Statement

The authors have declared no competing interest.

